# High-resolution tracking of microbial colonization in Fecal Microbiota Transplantation experiments via metagenome-assembled genomes

**DOI:** 10.1101/090993

**Authors:** Sonny TM Lee, Stacy A. Kahn, Tom O. Delmont, Nathaniel J. Hubert, Hilary G. Morrison, Dionysios A. Antonopoulos, David T. Rubin, A. Murat Eren

## Abstract

Fecal microbiota transplantation (FMT) is an effective treatment for recurrent *Clostridium difficile* infection and shows promise for treating other medical conditions associated with intestinal dysbioses. However, we lack a sufficient understanding of which microbial populations successfully colonize the recipient gut, and the widely used approaches to study the microbial ecology of FMT experiments fail to provide enough resolution to identify populations that are likely responsible for FMT-derived benefits. Here we used shotgun metagenomics to reconstruct 97 metagenome-assembled genomes (MAGs) from fecal samples of a single donor and followed their distribution in two FMT recipients to identify microbial populations with different colonization properties. Our analysis of the occurrence and distribution patterns post-FMT revealed that 22% of the MAGs transferred from the donor to both recipients and remained abundant in their guts for at least eight weeks. Most MAGs that successfully colonized the recipient gut belonged to the order Bacteroidales. The vast majority of those that lacked evidence of colonization belonged to the order Clostridiales and colonization success was negatively correlated with the number of genes related to sporulation. Although our dataset showed a link between taxonomy and the ability of a MAG to colonize the recipient gut, we also identified MAGs with different colonization properties that belong to the same taxon, highlighting the importance of genome-resolved approaches to explore the functional basis of colonization and to identify targets for cultivation, hypothesis generation, and testing in model systems for mechanistic insights.

## Background

Fecal microbiota transplantation (FMT), transferring fecal material from a healthy donor to a recipient, has gained recognition as an effective and relatively safe treatment for recurrent or refractory *Clostridium difficile* infection (CDI) [1–8]. Its success in treating CDI sparked interest in investigating FMT as a treatment for other medical conditions associated with intestinal dysbiosis, such as ulcerative colitis [9–11], Crohn’s disease (CD) [12–14], irritable bowel syndrome (IBS) [15,16]; and others, including metabolic syndrome [17], neurodevelopmental [18], and autoimmune disorders [19]. Despite the excitement due to its therapeutic potential, FMT also presents challenges for researchers and clinicians with potential adverse outcomes, including the transfer of infectious organisms [20] or contaminants from the environment [21,22]. A complete understanding of FMT from a basic science perspective is still lacking, as we have yet to determine the key microbial populations that are responsible for beneficial outcomes, as well as adverse effects.

Recent advances in high-throughput sequencing technologies, molecular approaches, and computation have dramatically increased our ability to investigate the ecology of microbial populations. Utilization of these advances at a proper level of resolution can lead to a better mechanistic understanding of FMT and identify new therapeutic opportunities or address potential risks. Most current studies on FMT use amplicons from marker genes, such as the 16S ribosomal RNA gene, to characterize the composition of microbial communities [23–26]. While providing valuable insights into the broad characteristics of FMTs, amplicons from the 16S ribosomal RNA gene do not offer the resolution to effectively identify populations that colonize recipients [27]. Other studies use shotgun metagenomics to annotate short reads and map them to reference genomes in order to track changes in the functional potential or membership in the gut microbial communities of recipients [28–30]. In a recent study, Li *et al.* [30] demonstrated the coexistence of donors’ and recipients’ gut microbes three months after FMT by mapping short metagenomic reads to reference genomes. Although this approach provides more information than marker gene amplicons alone, it is subject to the limitations and biases of reference genomic databases, is unable to characterize populations that do not have closely related culture representatives, and does not provide direct access to the genomic context of relevant populations for more targeted follow-up studies.

Metagenomic assembly and binning [31,32] is an alternative approach to characterizing microbial communities through marker gene amplicons or reference genomes. Here we used the state-of-the-art metagenomic assembly and binning strategies to reconstruct microbial population genomes directly from a single FMT donor, and tracked the occurrence of resulting metagenome-assembled genomes (MAGs) in two FMT recipients up to eight weeks.

## Methods

### Sample collection, preparation, and sequencing

We collected a total of 10 fecal samples; four samples from a single donor ‘D’ (a 30 year old male), and three samples from each of the two recipients ‘R01’ (a 23 year old male), and ‘R02’ (a 32 year old female) before and after FMT. Recipient samples originated from time points pre-FMT, four weeks after FMT, and eight weeks after FMT, while four samples from the donor were collected on four separate days two weeks prior to the transplantation. Both recipients had mild/moderate ulcerative colitis, had no genetic relationship to the donor, and received FMT through a single colonoscopy. We processed and stored all samples at -80°C until DNA extraction. We extracted the genomic DNA from frozen samples according to the centrifugation protocol outlined in MoBio PowerSoil kit with the following modifications: cell lysis was performed using a GenoGrinder to physically lyse the samples in the MoBio Bead Plates and Solution (5 – 10 mins). After final precipitation, the DNA samples were resuspended in TE buffer and stored at -20°C until further analysis. We prepared our shotgun metagenomic libraries with OVATION Ultralow protocol (NuGen) and used an Illumina NextSeq 500 platform to generate 2x150 nt paired-end sequencing reads.

### Metagenomic assembly and binning

We removed the low-quality reads from the raw sequencing results using the program ‘iu-filter-quality-minoche’ in illumina-utils [33] (available from https://github.com/merenlab/illumina-utils) according to Minoche *et al.* [34]. We then co-assembled reads from the donor samples using MEGAHIT v1.0.6 [35], used Centrifuge v1.0.2-beta [36] to remove contigs that are matching to human genome, and mapped short reads from each recipient and donor sample to the remaining contigs using Bowtie2 v2.0.5 [37]. We then used anvi’o v2.1.0 (available from http://merenlab.org/software/anvio) to profile mapping results, finalize genomic bins, and visualize results following the workflow outlined in Eren *et al.* [38]. Briefly, (1) the program ‘anvi-gen-contigs-database’ profiled our contigs using Prodigal v2.6.3 [39] with default settings to identify open reading frames, and HMMER [40] to identify matching genes in our contigs to bacterial [41] and archaeal [42] single-copy core gene collections, (2) ‘anvi-init-bam’ converted mapping results into BAM files, (3) ‘anvi-profile’ processed each BAM file to estimate the coverage and detection statistics of each contig using samtools [43], and finally (4) ‘anvi-merge’ combined profiles from each sample to create a merged anvi’o profile for our dataset. We used ‘anvi-cluster-with-concoct’ for the initial binning of contigs using CONCOCT [44] by constraining the number of clusters to 10 (‘–num-clusters 10’) to minimize ‘fragmentation error’ (where multiple bins describe one population). We then interactively refined each CONCOCT bin exhibiting conflation error (where one bin describes multiple populations) using ‘anvi-refine’ based on tetra-nucleotide frequency, taxonomy, mean coverage, and completion and redundancy estimates based on bacterial and archaeal single-copy genes. We classified a given genome bin as a ‘metagenome-assembled genomes’ (MAGs) if it was more than 70% complete or larger than 2 Mbp, and its redundancy was estimated to be less than 10%. We used ‘anvi-interactive’ to visualize the distribution of our bins across samples and ‘anvi-summarize’ to generate static HTML output for binning results. We further used CheckM v1.0.7 [45] to assess the completion and contamination of all bins and to assign taxonomy and used RAST [46] to ascribe functions to our MAGs. So that our analyses were not limited to the assembled portion of the data, we employed MetaPhlAn [47] to obtain the taxonomic community profiles in each sample from all short reads.

### Criteria for detection and colonization of MAGs

For each genome bin, anvi’o reports the percentage of nucleotide positions in all contigs that are covered by at least one short read based on mapping results, which is termed ‘portion-covered’. This statistic gives an estimate of ‘detection’ regardless of the coverage of a given genome bin. We required the portion-covered statistic of a genome bin to be at least 25% to consider it detected in a given sample. This prevented inflated detection rates due to non-specific mapping, which is not uncommon due to relatively well-conserved genes across gut populations. Finally, we conservatively decided that a MAG was transferred from the donor and colonized a given recipient successfully only if (1) it was detected in both samples that were collected from a the recipient at four and eight weeks after the FMT and (2) it was not detected in the pre-FMT sample from the same recipient.

### Statistical analyses

We performed cluster analyses on distribution profiles of MAGs and MetaPhlAn taxa using the R library vegan with Bray-Curtis distances of normalized values. We used the PERMANOVA (R adonis vegan) [48] test to measure the degree of similarity of the bacterial communities between the samples in the study. We further used similarity index (SIMPER) analysis to identify the taxa that contributed the highest dissimilarity between the samples. We classified the MAGs into four main groups based on their colonization characteristics in the recipients. We then performed a pairwise *t*-test (STAMP) [49] to ascertain any significant differences in the functional potential between the groups and carried out canonical correspondence analysis based on functional potential and the MAGs’ colonization characteristics.

### Data availability

Anvi’o profiles to reproduce all findings and visualizations in this study, as well as FASTA files and distribution statistics for each MAG, are stored under doi:10.5281/zenodo.185393. Raw metagenomic reads are also stored at the NCBI Sequence Read Archive under the accession number SRP093449.

## Results

The shotgun sequencing of genomic DNA from 10 fecal samples resulted in a total of 269,144,211 quality-filtered 2x150 paired-end metagenomic reads (Table S1). By co-assembling the donor samples, which corresponded to 115,037,928 of the quality-filtered reads, we recovered 51,063 contigs that were longer than 2.5 kbp and organized them into 444 genomic bins comprising a total of 442.64 Mbp at various levels of completion (Figure S1, Table S1). Using completion and size criteria, we designated 97 of our genomic bins as metagenome-assembled genomes (MAGs) (Figure 1, Table S1). Four major patterns emerged from the distribution of MAGs across individuals: MAGs that colonized both recipients R01 and R02 (Group I, n=22), MAGs that colonized only R01 (Group II, n=11), only R02 (Group III, n=8) and MAGs that did not colonize either of the recipients (Group IV, n=14) (Figure 1). We found no correlation between the abundances of MAGs in donor samples and their success at colonizing recipients (ANOVA, F=0.717, *p*=0.543). Table S1 reports the detection and mean coverage statistics for each MAG in each group.

**Figure 1.**
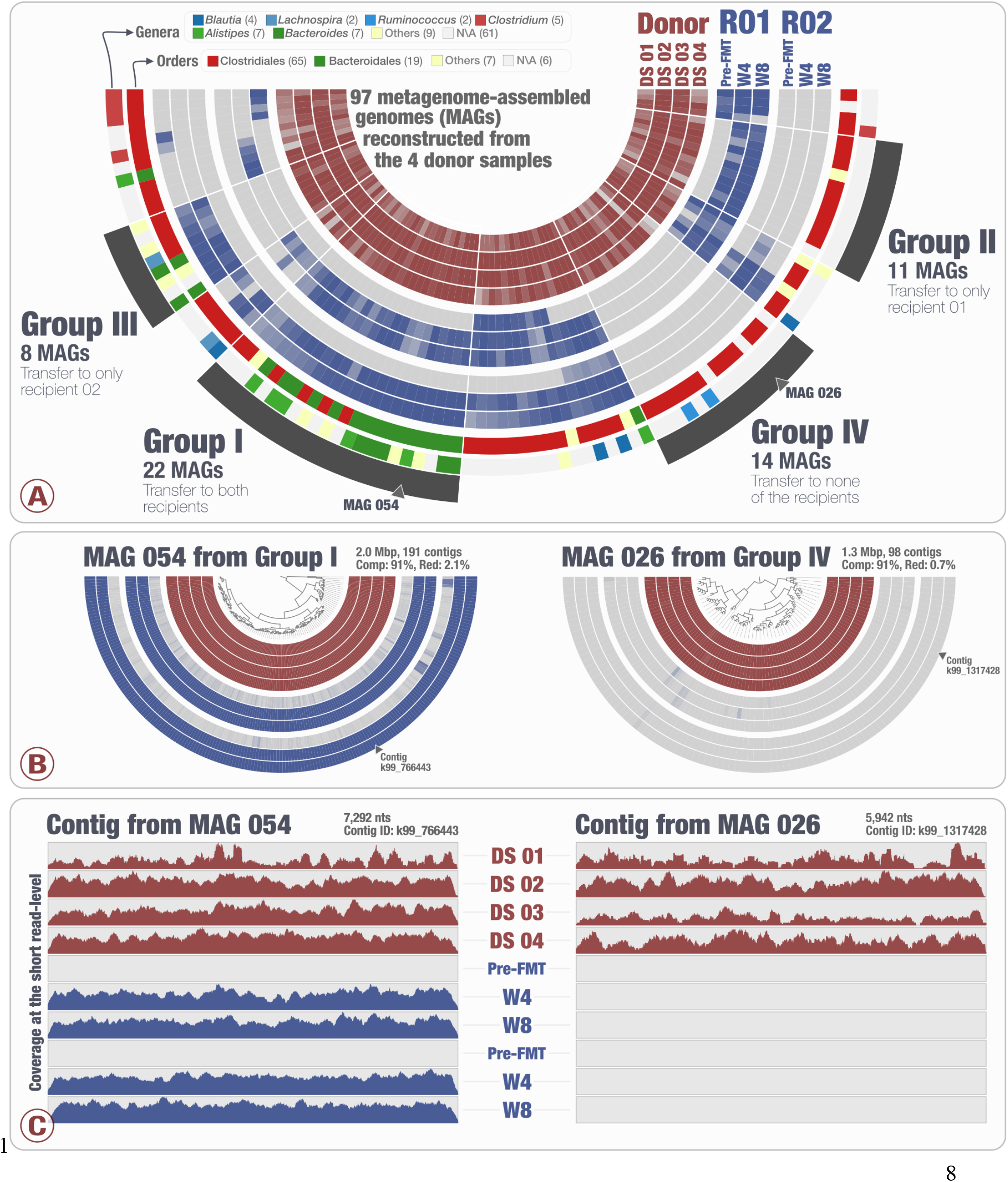
Distribution of MAGs across samples. Panel A shows the 97 MAGs and their level of detection in four donor samples (four inner circles) and in two recipients (R01 and R02) before FMT (pre-FMT), four weeks after FMT (W4), and eight weeks after FMT (W8). Bars in the layers that represent donor and recipient samples indicate the level of detection of a given MAG in a given sample. The outermost two circles display the genus- and order-level taxonomy for each MAG. Panel A also displays the selection of four groups: Group 1 with 22 MAGs that colonized both recipients, Group II with 11 MAGs that colonized only R01, Group III with 8 MAGs that colonized only R02, and finally Group IV with 14 MAGs that colonized neither recipient. Panel A displays the average detection of each MAG and Panel B displays the coherence of detection for each contig in two example MAGs. Panel C displays the coherence of detection for each nucleotide positions in two example contigs from the MAGs displayed in Panel B.

The taxonomy of 15 of the 22 MAGs that colonized both recipients resolved to the order Bacteroidales (Figure 1). Besides Bacteroidales, Group I also included six MAGs that were classified as order Clostridiales and one MAG as Coriobacteriales. CheckM partitioned the Group I MAGs into two genera, *Bacteroides* (n=5) and *Alistipes* (n=5). Eight MAGs in this group were not assigned to a specific genus. In contrast to the Bacteroidales-dominated Group I, of the 14 MAGs that did not colonize recipients (Group IV) resolved to the order Clostridiales. The remaining three MAGs were not assigned any taxonomy at the order level. The only genus-level annotation for the MAGs in Group IV was *Ruminococcus* (n=2). Overall, CheckM did not assign any genus-level taxonomy to 20 of the 36 MAGs that colonized either both recipients (Group I) or none (Group IV).

MAGs that colonized only one recipient’s gut did not show a consistent taxonomic signal. While 9 of 11 MAGs that colonized only R01 (Group II) were assigned to the order Clostridiales, only 4 of 8 MAGs that colonized R02 (Group III) were assigned to that order (Figure 1, Table S1). The remaining MAGs in Group III were assigned to Bacteroidales (n=2), Burkholderiales (n=1), or not assigned (n=1).

We used non-metric multidimensional scaling (nMDS; 2D Stress: 0.03 with Bray-Curtis similarity index) on square-root normalized values of the microbial community profiles based on the average coverage of the 97 MAGs as well as the genus-level taxonomy as characterized by MetaphlAn using all metagenomic short reads. Both analyses revealed an increased similarity between the donor microbiota and the recipients following the FMT experiment (Figure 2). The donor and recipient bacterial community profiles differed significantly from each other before FMT (PERMANOVA, pseudo-F=11.952, *p*=0.002; Figure 2) and bacterial community profiles within each recipient shifted significantly after FMT (PERMANOVA, pseudo-F=3.993, *p*=0.026; Figure 2). Based on metagenomic short reads, the microbial community structure in both R01 and R02 were more than 60% similar to the donor microbiota after FMT (Figure 2). Furthermore, similarity percentage analysis (SIMPER) of the community structure based on genus-level taxonomy suggested that the two recipients were 61.24% similar after FMT and that *Bacteroides* was responsible for the largest fraction (14.65%) of the recipient sample differences between pre-FMT and four weeks after FMT. There were no significant changes in the recipients’ bacterial community between week four and eight post-FMT (PERMANOVA, pseudo-F=0.223, *p*=0.665; Figure 2).

**Figure 2.**
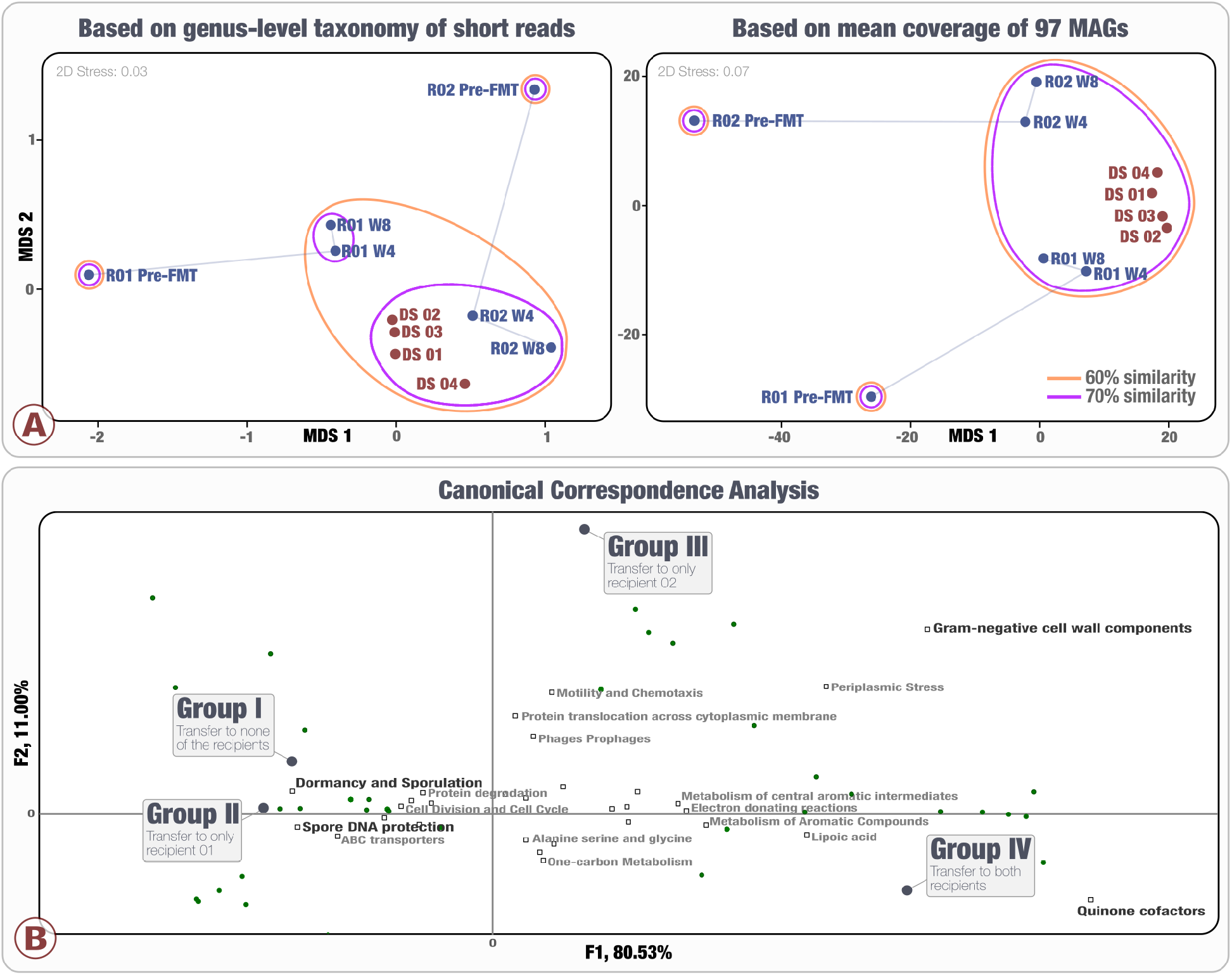
(A) Non-metric multidimensional scaling based on microbial community profiles at the genus level of short reads annotated by MetaPhlAn and based on mean coverage of 97 MAGs. Clustering employed average linkage with Bray-Curtis similarity index on square-root normalized values. Labels represent the donor (D) with four sample replicates (S01 – S04), and recipients (R01, R02) before FMT (Pre-FMT), four weeks (W4) and eight weeks after FMT (W8). (B) Canonical correspondence analysis of 97 MAGs based on the 29 significant functional subcategories and the detection of donor’s microbiota in the recipients.

To investigate whether there was a functional link between MAGs and their success of colonization, we studied 500 functions and 110 sub-systems assigned by RAST across our 97 MAGs (Table S2). We performed a canonical correspondence analysis (CCA) to determine whether functional markers could be used as an indicator for groups of bacteria that were more or less likely to colonize recipients. CCA (pseudo-F=1.746, *p<*0.0001) revealed that the MAGs that colonized both recipients (Group I) possessed a higher relative abundance of genes coding for quinone cofactors. Group I also showed potential functions involving gram-negative cell wall components, periplasmic stress, and metabolism of aromatic compounds and their intermediates. In contrast, the MAGs that did not colonize any of the recipients carried higher number of genes related to dormancy and sporulation, spore DNA protection, and motility and chemotaxis (Figure 2, Table S2).

## Discussion

Our study demonstrates that genome-resolved metagenomics can facilitate high-resolution tracking of the donor populations in recipient guts after FMT experiments by revealing bacterial populations with differential colonization properties. Previous studies reported an increase in relative abundance of *Alistipes* [23,24,50–52] and *Bacteroides* populations after FMT experiments [23–26,30]. The success of the order Bacteroidales was also striking in our dataset: 15 of the 19 Bacteroidales MAGs we identified in the donor successfully colonized both recipient guts (Figure 1). Although taxonomic signal was relatively strong, our results also showed that taxonomy is not the sole predictor of transfer, as MAGs that resolved to the same genera (i.e., *Alistipes*, *Bacteroides*, and *Clostridium*) showed different colonization properties. In addition, taxonomic annotation of a large fraction of MAGs in our study did not resolve to a genus name, which suggests that bacterial populations that have not yet been characterized in culture collections may be playing important roles in FMT treatments.

Although a substantial number of studies report successful medical outcomes of FMT experiments [3,7,53,54], a complete understanding of this procedure from the perspective of microbial ecology is still lacking. Studying FMT as an ecological event, and the identification of the key fecal components that facilitate the procedure’s success as a treatment for intestinal disorders require the characterization of the transferred microbial populations at a high level of resolution. In contrast to operational taxonomic units identified through 16S rRNA gene amplicons that often combine multiple populations into a single unit [55,56], MAGs reconstructed directly from the donor samples can provide enough resolution to guide cultivation efforts. A recent effort by Vineis et al. [57] demonstrated this principle by first identifying populations of interest using MAGs reconstructed from a gut metagenome and then using the genomic context of those MAGs to screen culture experiments from the same gut sample to bring the target population to the bench. A similar approach in the context of FMTs can provide opportunities to design experiments to explore the functional basis of colonization in controlled systems.

The complete transfer of fecal matter between individuals comes with various risks. For instance, a recent meta-analysis of 50 peer-reviewed FMT case reports reported 38 potentially transfer-related adverse effects in FMT patients in 35 studies, including fever, sore throat, vomiting, abdominal pain, bowel perforation, rhinorrhea, transient relapse of UC and CDI, and in one case, death, due to temporary systemic immune response to the applied bacteria [58]. Besides bacteria, FMT can transfer viruses, archaea, and fungi, as well as other agents of the donor host such as colonocytes [59], which may affect the recipient’s biology in unexpected ways. A more complete understanding of the microbial ecology of FMTs would identify precisely what needs to be transferred, so that recipients benefit from the positive outcomes of FMT without incurring medical risks from uncharacterized biological material.

A recent study by Khanna *et al.* [60] reported high rates of success with the treatment of patients with primary *Clostridium difficile* infection (CDI) using an investigational oral microbiome therapeutic, SER-109, which contains bacterial spores enriched and purified from healthy donors. However, Seres Therapeutics announced more recently that interim findings from the mid-stage clinical study of SER-109 failed to meet their primary goal of reducing the risk of recurrence for up to eight weeks [61]. In our study, the MAGs that failed to colonize any of the recipients were significantly enriched for spore-formation genes. Interestingly, Nayfach et al. [62] recently made a similar observation regarding the transmission of bacteria and sporulation in a different system, vertical transmission between mothers and their infants. Populations with high vertical transmission rates had lower number of genes related to sporulation [62]. These observations suggest that dismissing non-spore forming bacteria may decrease the efficacy of FMT therapies due to limited colonization efficiency, and deeper insights into the functional basis of microbial colonization warrants further study.

Identifying and using bacterial populations associated with positive health outcomes and that harbor high colonization properties may result in more effective therapies compared to cleansing all but spore-forming bacteria to avoid the transfer of pathogens. The analytical strategy adopted in our study can facilitate the identification of bacterial population genomes that may be critical to the success of FMT due to their colonization properties, and provide genomic insights to leverage our investigations beyond associations, and ultimately reveal the mechanistic underpinnings of this procedure.

## Declarations

### Ethics approval and consent to participate

The study was reviewed and approved by the University of Chicago Ethics Committee and by the University of Chicago Institutional Review Board (IRB 132-0212). Written and informed consent was obtained for all participants.

### Consent for publication

Not applicable

### Availability of data and materials

All data generated and analyzed during this study are included in this published article and its supplementary information files.

### Authors’ contributions

SAK, NJH, DAA, and DTR designed the study, collected, and processed the patient samples. STML, TOD, HGM, and AME generated, processed, and analyzed the sequencing data. STML, and AME wrote the manuscript. All authors read and approved the final manuscript.

